# A tunable affinity fusion tag for protein self-assembly

**DOI:** 10.1101/2025.01.14.633037

**Authors:** Hannah Kimbrough, Jacob Jensen, Tayla Miller, Jeffrey J. Lange, Randal Halfmann

## Abstract

The concentrations of individual proteins vary between cells, both developmentally and stochastically. The functional consequences of this variation remain largely unexplored due to limited experimental tools to manipulate the relationship of protein concentration to activity. Here, we introduce a genetically encoded tool based on a tunable amyloid that enables precise control of protein concentration thresholds in cells. By systematically screening dipeptide repeats, we identified poly-threonine alanine (poly-TA) as an ideal candidate due to its unique ability to form amyloid-like assemblies with a negligible nucleation barrier at arbitrarily chosen concentration thresholds. We demonstrate that the saturating concentration (C_sat_) of poly-TA can be finely tuned by adjusting the length of uninterrupted TA repeats, even while maintaining length and composition, providing a modular system for manipulating protein solubility. This tool offers a powerful approach to investigate the relationship between protein concentration, phase separation, and cellular function, with potential applications in cell-, developmental-, and synthetic biology.

## Introduction

Cell-to-cell differences in protein concentration underpin biology. Genetically programmed differences in the concentrations of certain transcription factors, metabolic enzymes, and signaling proteins functionally diversify every cell type in our bodies, while stochastic differences in protein concentration facilitate adaptation and the evolution of microbes and tumors (1–4).

Our ability to observe and measure such differences have far outpaced our ability to manipulate them. Single cell omics approaches are rapidly uncovering previously hidden diversity in genetically identical cell populations. This diversity typically involves *quantitative* differences in the measured (or inferred) levels of specific proteins, and these differences are presumed to translate into different cellular phenotypes and functions. This presumption will be flawed for some proteins, however, because the relationship of activity to concentration differs between proteins. For some, activity will increase monotonically with concentration. For others, it will increase sharply at a specific concentration dictated by cooperative self-assembly. For still others, it will be buffered against concentration via phase separation (5) or even fall with concentration as ordered aggregates nucleate and sequester the protein (6). Hence, we cannot infer protein activity or phenotypic consequence from expression alone. Unfortunately, we lack the ability to rigorously interrogate and manipulate the relationship of protein concentration to function in cells, and therefore the ability to fully understand the relationship of cell diversity to organismal function and evolution.

Progress toward this goal would be aided by a genetically encodable device whose activity changes from one state to another as its concentration passes an experimentally defined arbitrary threshold. This device would offer an advantage over existing tools to manipulate protein activity levels, such as inducible promoters and protein degradation tags (degrons), which respond to input from the experimentalist rather than the biological system itself. This device would be “smart” -- responding directly to its own concentration in the cell, while ignoring other proteins and variables like pH and temperature. This would allow the experimentalist to impose a lower or upper bound on the concentration at which it activates, and to program cells to respond autonomously when that concentration threshold is subsequently breached. Coupling this device to various effectors would enable biologists to manipulate cell diversity with previously unachievable precision. For example, it could be coupled to a degron to cap the concentration of a protein to which it is fused, or coupled to a caspase to selectively kill cells with slightly higher expression of an oncogene, or delete cell types according to small differences in marker protein expression, allowing for the functional interrogation of that cell type in the organism.

Protein phase separation provides the mechanistic basis for such a tool. Specifically, phase separation involves the highly cooperative interaction of multivalent protein molecules whose effective concentration exceeds a threshold known as the saturating concentration (C_sat_). The outcome of phase separation is two discrete regions of space with separate, fixed concentrations of the protein -- the lower concentration solution phase wherein the proteins do not interact, and the higher concentration condensed phase, or condensate, wherein the proteins interact continuously. Condensation can involve only a change in density, resulting in liquid-liquid phase separation (LLPS), or it can be coupled with the acquisition of translational and orientational order, resulting in crystallization. The activity of proteins differs quantitatively or qualitatively between the two phases. Numerous synthetic liquid condensates have been designed to exploit this principle by manipulating effective concentrations or valencies of the proteins with light, temperature, or small molecules (7). The multivalent interaction moieties range from charged or aromatic residues, to coiled-coils, to entire folded domains, all joined by disordered linkers for a total polypeptide length exceeding hundreds of residues (8,9). Unfortunately, these long lengths limit the utility of such condensates as fusion partners in the cell. Long proteins are costly (both to the experimentalist and to the cell), have increased potential to sterically interfere with protein activity, and limit the sensitivity of payload proteins by competing for physical volume in the condensate. Intrinsically disordered proteins that form the basis of many liquid condensates also tend to interact promiscuously with other cellular components, which alters cell function (10) and renders them susceptible to differences in cell properties other than their own concentration (11–14). In addition, the sequence complexity of natural condensate-forming proteins creates multiple energy scales for self-interaction that dampens their responsiveness to concentration thresholds (15,16).

Synthetic crystals bypass some of the limitations of LLPS by requiring a specific three-dimensional geometry that enhances specificity, and many have been developed as materials and biosensors (17,18). Unfortunately, however, folded crystallizing domains also require relatively long encoding sequences, and their affinity cannot be tuned by simply lengthening the sequence as for LLPS.

A specific kind of condensate could theoretically merge the useful properties of liquids and globular crystals, while avoiding the limitations. Amyloids are quasi one-dimensional crystals defined by translational ordering of polypeptide backbones rather than a derived three-dimensional fold. This means that affinity can in some cases be tuned by simply lengthening the sequence (19,20), as in liquid condensates. Amyloids nevertheless achieve extraordinary self-selectivity and stability even from short sequences, owing to a continuous array of repeating hydrogen bonds and interdigitated side chains along their axes (21). Amyloids tend to form from intrinsically disordered sequences whose backbones are accessible for intermolecular beta sheet formation. Unfortunately, this imposes a large entropic cost for amyloid formation, which manifests as a nucleation barrier that tends to delay assembly until the protein’s concentration far exceeds its C_sat_. This clearly presents a challenge for using amyloids as concentration-sensing devices.

To overcome the many challenges to exploiting phase separation to sense and respond to protein concentration, we here identify and characterize a simple and short amyloid-forming sequence that can be easily tuned to assemble with a negligible nucleation barrier at arbitrary thresholds of concentration. Our tunable concentration-sensing tag is nontoxic, modular, and insensitive to the cellular environment. We anticipate future demonstrations of its use as a genetic fusion to diverse proteins to control and manipulate cell state.

## Results

The universe of amyloid-forming sequences is vast (22). Our simple goal to tune affinity through small variations in peptide length allows us to consider only a tiny fraction of that universe -- simple tandemly repeated sequences. We can further focus our search to dipeptide repeats by considering that the core structural element of amyloid -- a pair of beta sheets interacting across a tight interface or steric zipper -- can be encoded at every other position in sequence. Indeed, this property was recently exploited in the form of YK dipeptide repeats to create an ATP-responsive synthetic amyloid in mammalian cells (19).

We observed that AlphaFold2 can accurately model functional (“native”) amyloids (**Figure S1A**), and therefore reasoned that it could usefully inform a search for amyloid-forming dipeptide repeats. We therefore used AlphaFold2 and a custom script to screen all 380 dipeptide repeat sequences for native-like amyloid formation. For each repeat sequence, we generated homopentamer models of peptides containing 20 repeats (40 residues), a length chosen to exceed the short cores of functional intracellular amyloids such as Ripk3 (19-22 residues; (23,24)) while remaining short enough to minimize monomeric structure. We anticipated that this short length would prevent interactions or steric interference with eventual fusion partners, while still being long enough to tune affinity by truncating the sequence (**Figure S1B**). Models were analyzed by metrics for confidence (predicted local distance difference test, pLDDT; **Figure 1A**) and amyloid-like features (□ structure, backbone planarity and interaction energy; **Figure 1B-C**). We found that the orientations of the two residues in the repeat unit did not strongly influence the models (e.g. (AY)40 scored similarly as (YA)40), as expected given that only the two terminal residues’ properties differ. We therefore considered them as the same sequence. We eliminated from consideration dipeptide repeats whose models all had low confidence or lacked amyloid-like structure. We further excluded one of the remaining candidates (VK) based on the sensitivity of polycations to endogenous anions (19,25) and their tendency to toxically sequester endogenous polyanions (26). This left just three dipeptides for further consideration: NA, TA, and YA (**Figure 1D**). Given that all three sequences feature alanine, we also included GA, an alanine-containing dipeptide repeat that was not identified as a candidate but that had nevertheless been shown to form amyloid (27).

**Figure 1.**
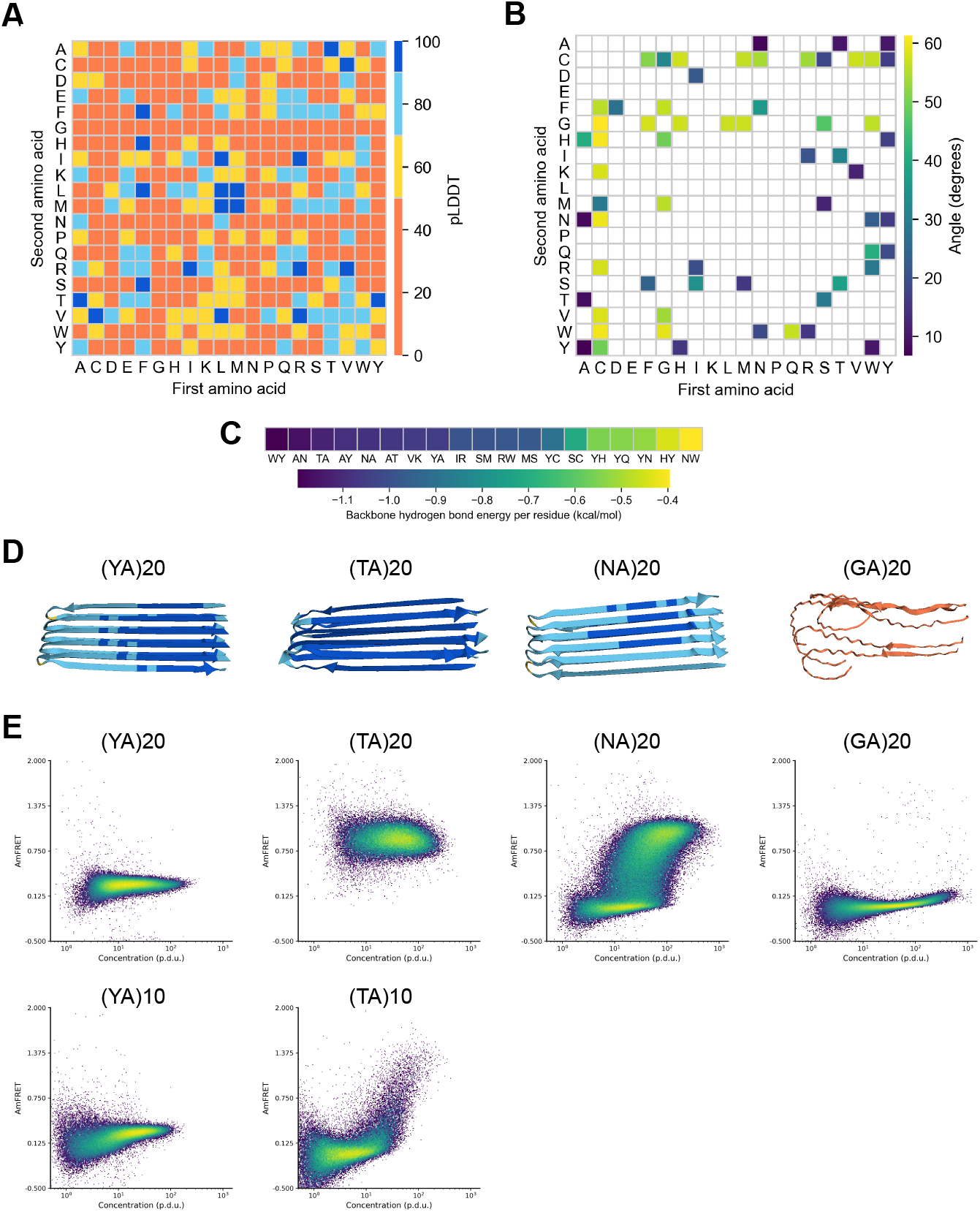
Systematic screen for native-like amyloid formation by dipeptide repeat sequences. (a) Heatmap of pLDDT of all top ranked AlphaFold2 models of homo-pentamers of dipeptide repeat (20x) polypeptides. (b) Heatmap of maximum average carbonyl angle of beta sheets in putative amyloid core containing structures. (c) Per residue backbone hydrogen bond energy of putative amyloid structures with maximum average carbonyl angle less than 20 degrees. (d) Top ranked AlphaFold2 models of candidate dipeptide repeats. (e) Representative DAmFRET plots of dipeptide sequences of length 40 (top) and 20 (bottom).

We proceeded to compare the amyloid propensities of these six sequences using distributed amphifluoric FRET (DAmFRET, (6)). Specifically, we expressed each sequence as a 40 residue-long peptide fused to the monomeric fluorescent protein, mEos3.1, in budding yeast cells. We then exposed the culture of cells to violet light to photoconvert an optimized fraction of mEos3.1 molecules from green to red, and then used flow cytometry to quantify ratiometric FRET between the two fluorescent forms (AmFRET) as a function of each protein’s intracellular concentration (quantified as the ratio of directly excited acceptor fluorescence intensity and side scatter (SSC), a proxy for cell volume (28)). The duration of protein expression (hours) in a DAmFRET experiment is long relative to timescales of LLPS, but short relative to timescales of typical amyloid nucleation in vivo (29,30). Consequently, for query proteins that condense with negligible nucleation barriers, such as by LLPS, cells uniformly acquire AmFRET beyond the protein’s C_sat_. In contrast, for query proteins that condense with large intrinsic (conformationally determined) nucleation barriers, such as for most amyloids, cells acquire AmFRET semistochastically to yield a discontinuous distribution. Hence, DAmFRET facilitates comparative assessments of sequence features that encode intrinsic nucleation barriers across proteins. The desired dipeptide sequence will produce a sharp and uniform gain of AmFRET beyond a threshold concentration, but lack AmFRET below that concentration.

The DAmFRET data confirmed robust self-assembly by all three of the sequences that scored well in the AlphaFold2 screen (**Figure 1E**): (NA)20, (TA)20, and (YA)20. All cells expressing (TA)20 and (YA)20 exhibited high AmFRET, indicating these sequences form stable self-assemblies that lack a detectable nucleation barrier across the range probed by DAmFRET (approximately 2 to 200 μM, (6)). Cells expressing (NA)40 exhibited a discontinuous jump in AmFRET over a range of concentrations, indicating a large intrinsic nucleation barrier typical of amyloid (but undesired for our application), and hence was not considered further. (GA)20, the sequence that had not scored well by AlphaFold2, did not appreciably assemble. To determine if the affinity of poly-TA or -YA assemblies can be tuned by varying their repeat numbers, we retested them as shorter peptides with only 10 repeats. At this length, poly-YA was still fully assembled at all expression levels, restricting its potential window for tunability. We therefore did not explore it further. In contrast, poly-TA now remained soluble until reaching very high concentrations. We proceeded to test a series of poly-TA variants from 8 to 20 repeats (**Figure 2A**). We found that C_sat_ fell with length across this range (**Figure 2B**). As an orthogonal measure of the stability of poly-TA assemblies, we analyzed their stabilities and length dispersions using semidenaturing detergent-agarose gel electrophoresis (SDD-AGE). (TA)20 formed detergent-resistant polydisperse species (**Figure S1C**) that are diagnostic of amyloid-like aggregation (31).

**Figure 2.**
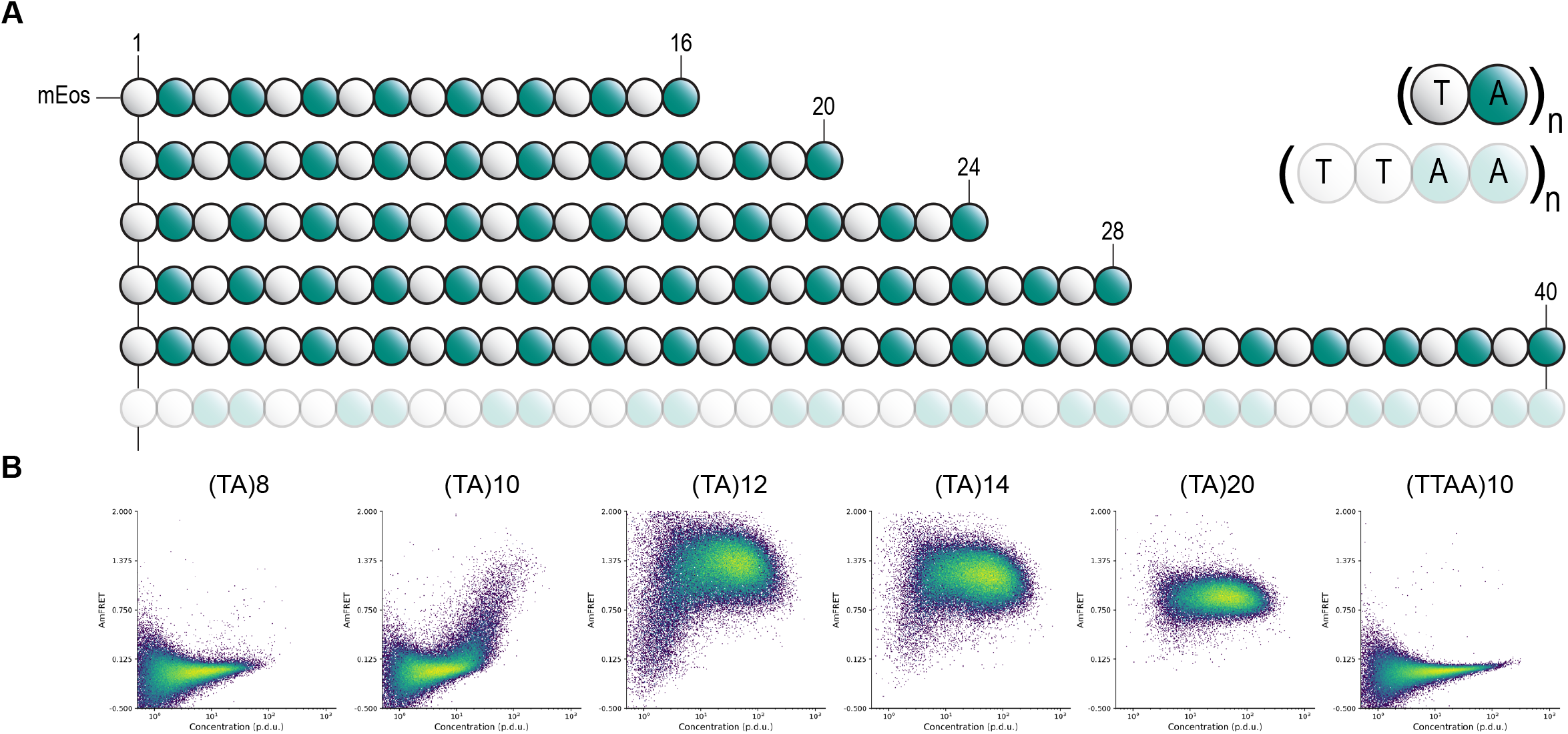
poly-TA length tunes saturating concentration. (a) TA dipeptide repeat length and orientation schematic and (b) representative DAmFRET plots.

The ideal tag for fusion protein self-assembly would allow experimentalists to precisely manipulate the threshold concentration for self-assembly without changing the length or sequence composition of the tag, thereby eliminating these variables that could potentially confound experiments. The relationship of dipeptide sequence to amyloid structure suggests a simple way to achieve this ideal -- by reversing the order of residues in some of the repeats. The AlphaFold2 model of poly-TA amyloid has the alanine residues forming a tightly interdigitated hydrophobic interface between two closely apposed sheets (**Figure S1E**). The hydrophilic and relatively bulky threonine residues cannot be accommodated in this interface and are strictly confined to the exterior surface. If this model is correct, then disrupting the repeating dipeptide pattern will disrupt self-assembly. We therefore reversed the order of T and A in every other repeat in the otherwise highly amyloidogenic (TA)20 repeat sequence, yielding a sequence of (TTAA)10. This sequence failed to form amyloid as modeled by AlphaFold2 (**Figure S1E**), and in agreement, was completely soluble at all expression levels when analyzed by DAmFRET and confocal microscopy (**Figure 2A, Figure S1D**). We conclude that TA is the most suitable dipeptide repeat for a tunable self-assembling tag.

Having confirmed the selectivity of the poly-TA steric zipper for alternating T and A residues, we next designed a series of 28-residue sequences all containing 14 Ts and 14 As, uniformly distributed but with different lengths of uninterrupted dipeptide repeat tracts (**Figure 3A**). Using DAmFRET, we observed a fine gradation of C_sat_ values over a greater-than tenfold range of concentrations (**Figure 3B**), and a tight correlation between C_sat_ and the uninterrupted tract length of each sequence (**Figure 3C**).

**Figure 3.**
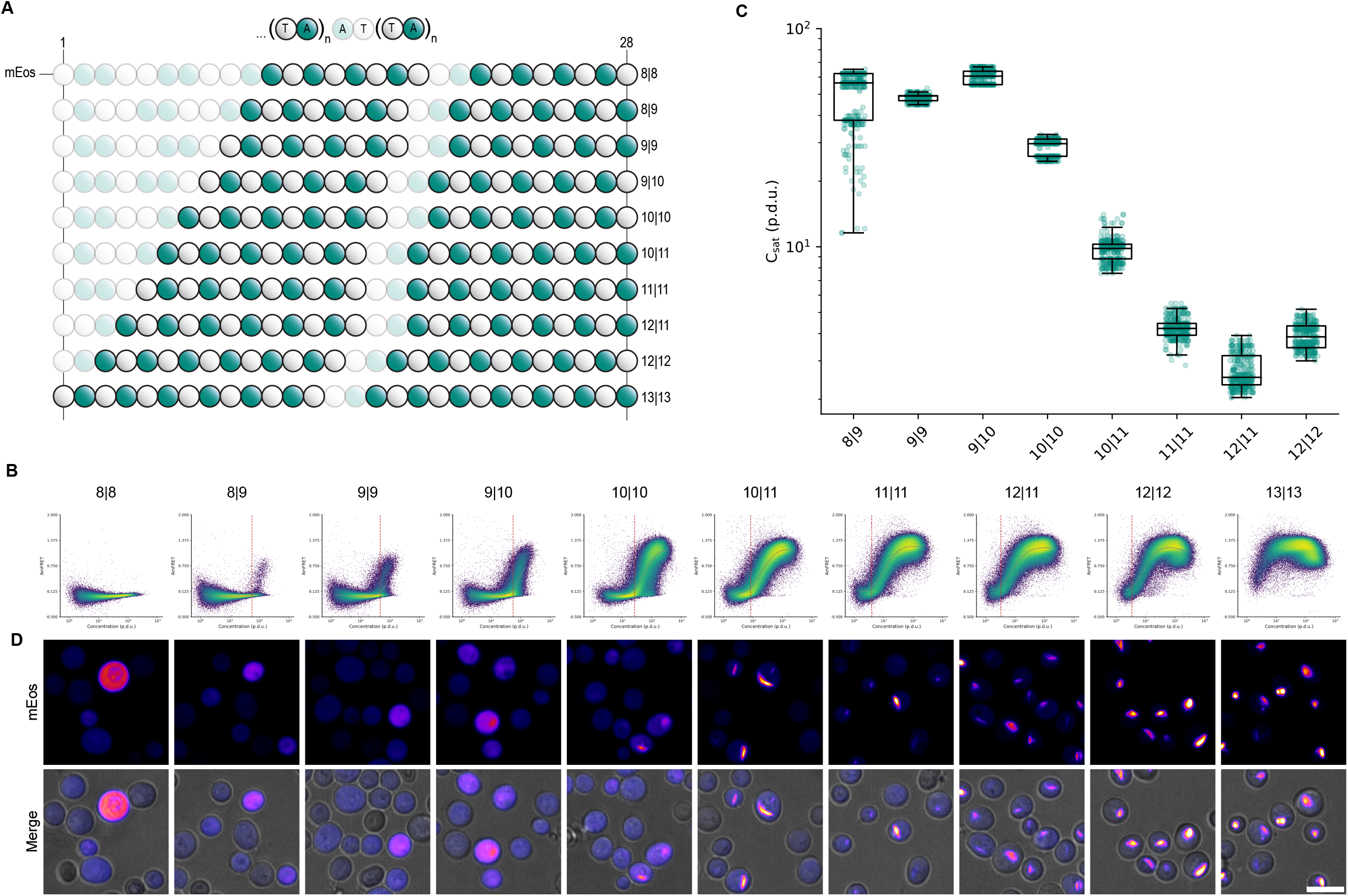
poly-TA affinity tunes saturating concentration. (a) TA dipeptide repeat tract length schematic. (b) Representative DAmFRET plots (top) fitted to spline function (gray) used to calculate the saturating concentration (red dotted line) and representative confocal microscopy images (bottom) of yeast cells expressing varying poly-TA tract lengths at a range of concentrations. poly-TA tract lengths of 8|8 and 13|13 were excluded from spline fitting analysis as cells either did not reach or exceeded, respectively, the saturating concentration detectable via DAmFRET. Scale bar, 10 μm. (c) Boxplot of saturating concentration of poly-TA tract lengths (n=300).

We next analyzed the intracellular distribution of the fusion proteins by confocal fluorescence microscopy. Consistent with the DAmFRET data, we observed that the proteins were fully diffuse at low concentrations and tract lengths, but became punctate as concentration and tract length increased (**Figure 3B**). The puncta were spindle-shaped at the lowest tract lengths supporting assembly, consistent with bundles of ordered filaments. They became rounder and less regular with increasing tract length, presumably reflecting a branched or gel-like ultrastructure as a consequence of their exceeding the optimum strand length of the amyloid core (32,33).

The existence of a threshold concentration for aggregation in cells implies that the cytoplasm has a finite carrying capacity for the protein. In principle this could result from saturation of macromolecular binding partners such as protein chaperones, which would preclude the protein’s use as an accurate sensor of its own concentration across biological systems. To account for this possibility, we tested the impact on C_sat_ of fusing poly-TA to solubility-enhancing tags. Two well-characterized tags in particular -- maltose binding protein (MBP) (34) and EB250 (35) -- have extremely different physico-chemical properties but nevertheless share a tendency to passively enhance the solubility of diverse proteins to which they are fused. We found that, as expected, both greatly increased (by more than tenfold) the C_sat_ of poly-TA. Because their fusion could not have changed the abundance of poly-TA relative to potential binding partners, we conclude that the apparent C_sat_ for poly-TA is an intrinsic property of its sequence in cytosol (**Figure 4A-B**). We note that this simple analysis could be easily extended to diverse fusion tags to rapidly and quantitatively compare their efficacies, or to screen libraries of candidate solubility enhancers.

**Figure 4.**
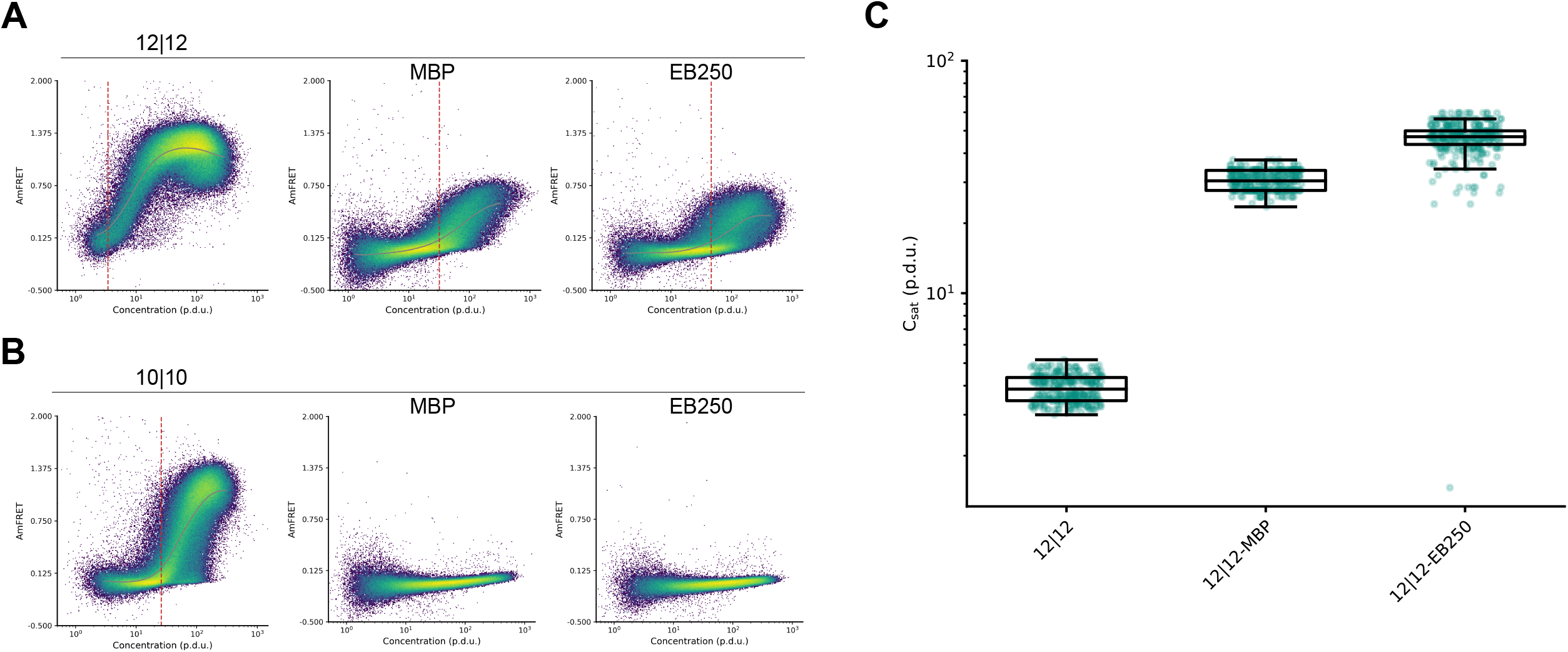
poly-TA fusion to solubility tags increases saturating concentration. (a) Representative DAmFRET plots of yeast cells expressing poly-TA peptide with total length of 28 residues with two continuous TA tracts of length 12 either independently (left), or fused to MBP (middle) or EB250 (right). (b) Representative DAmFRET plots of yeast cells expressing poly-TA peptide with total length of 28 residues with two continuous TA tracts of length 10 either independently (left), or fused to MBP (middle) or EB250 (right). DAmFRET plots are fitted to a spline function (gray) and used to calculate saturating concentration (red dotted line). Poly-TA fusions that were completely monomeric were excluded from spline fitting analysis as cells did not reach the saturating concentration required for assembly. (c) Boxplot of saturating concentration of poly-TA tract length 12 with solubility tag fusions (n=300).

## Discussion

We recognized that biologists presently lack a facile way to rigorously interrogate and manipulate the relationship of protein concentration to function in cells. We sought to address that need, in part, by exploiting the properties of phase separation to sense and respond to protein concentration with an arbitrary and tunable threshold, while exploiting the properties of amyloids to increase the specificity and resilience of phase separation to varied biological contexts. Doing so necessitated that we restrict the intrinsic kinetic barrier to self-assembly that tends to make amyloid formation relatively insensitive to concentration (6,36). The result of this undertaking -- a dipeptide (TA) repeat of varying length -- achieves all objectives. We are now poised to couple this module with proximity-dependent effector modules, such as caspases (37), to link concentration-sensing to desired cellular outcomes.

Our effort to deploy an amyloid as an experimental tool for cell- and developmental biology may seem misguided given the notorious relationship of amyloids to disease. We believe otherwise. A rapidly growing list of amyloids are now appreciated to serve essential biological functions (38,39). The toxicity formerly attributed to pathologic amyloids themselves is now more closely attributed to their process of formation, which drives an accumulation of noxious metastable oligomers (40,41). The folding pathway for functional amyloid sequences tends to bypass such species. With its exceptionally small intrinsic nucleation barrier that precludes supersaturation, poly-TA almost certainly mimics the latter. With its low hydrophobicity and absence of charges, we further expect it to be relatively insensitive to varying intracellular conditions such as pH, temperature, and metabolites. Future investigations will explore the uniformity of C_sat_ across biological systems and whether it is indeed biologically inert.

Although we identified poly-TA from an empirical survey of dipeptide repeats, we might as well have identified it from purely theoretical considerations of sequence-structure relationships. Alanine has the smallest sidechain of any residue (other than the hydrogen “side chain” of glycine) and consequently has the lowest entropic penalty for side chain interdigitation in the steric zipper core of amyloid fibrils. Threonine has a higher propensity for □-structure than any other hydrophilic residue (42). It is therefore perhaps unsurprising that a repeated alternation of these two residues would allow for native-like folding into amyloid fibrils.

Poly-TA had previously been characterized as an artificial mimetic of CAT tails, C-terminal extensions of alanine and threonine residues appended to stalled nascent polypeptides by ribosome quality control machinery (43). Although endogenous CAT tails are now appreciated to lack regular sequence, this line of inquiry led to the recent discovery that poly-TA can form amyloid as a consequence of its alternating hydrophilic-hydrophobic pattern (44–46). Its tunability and unusual kinetic properties had not previously been investigated, however.

The direct relationship of poly-TA tract length to solubility shown here provides a simple framework for experimentalists to precisely manipulate the solubility of their protein of interest by genetically fusing it to TA tracts of the desired length. Indeed, in work to be published separately via collaboration, we deploy poly-TA as a modular and tunable self-assembling module in the context of human innate immune signaling (47). Specifically, poly-TA was used to reconstitute the scaffolding property of death domains in the MyDDosome, a membrane-localized signaling complex that forms in response to certain cytokines and pathogen-derived ligands. By varying the number of TA repeats in a chimeric MyDDosome, Lichtenstein et al. demonstrated that signaling only occurred above a threshold of 15 repeats, whereupon increasing the number of repeats increased both the stability of the complex and its signaling output in response to cytokine. Hence, the opportunity afforded by poly-TA to tune the stability of the chimeric fusion protein’s self-assembly allowed Lichtenstein et al. to confirm their hypothesis that polymer stability governs signalosome functionality. We anticipate poly-TA will find many similar applications moving forward.

## Materials and Methods

### Plasmid and yeast strain construction

*Saccharomyces cerevisiae* strains rhy2145, rhy3082, and rhy3279 were previously described (30,48). All strains are [*pin*^−^] and isogenic with the exception of mating type and an integration of BDFP1.6:1.6 at the *PGK1* locus (in rhy3082). Plasmids (**Table S1**) were constructed by replacing the insert in vector V12 (6) with yeast codon-optimized fragments (Genscript) to create open reading frames with mEos3.1 followed by an alpha helical linker and the repeat sequence of interest. Plasmids expressing solubility tags N-terminally fused to mEos3.1 were constructed by inserting yeast codon optimized MBP or EB250 into rhx5422 and x5425. Yeast strains were transformed using a standard lithium acetate procedure.

### AlphaFold structural predictions and analysis

All dipeptide repeat combinations at a total peptide length of 40 amino acids were modeled as pentamers using AlphaFold2 (49) as implemented in ColabFold 1.5.1 (50) as the “alphafold2_ptm” model. No additional sequences were included in the multiple sequence alignments, and ColabFold was run with 25 recycles. The 5 models for each dipeptide repeat were sorted by the “multimer” rank metric, and the highest ranking models were analyzed further. Secondary structure labels were assigned to residues using DSSP (51). Models were considered to contain a putative amyloid core if there were at least 2 beta sheets conserved at the same residue positions across at least 3 chains, with at most 7 non-beta strand residues between the beta sheets. To assist in measuring planarity of the putative amyloid cores, vectors were calculated from the C atom to O atom of the backbone of each residue. Within each beta sheet having at least 4 residues per strand, the average angle between these vectors compared to the vector of the middle residue listed for that sheet (typically at the center of the strand for that sheet within chain C) was calculated. Angles above 90 degrees were subtracted from 180 to account for the alternating directions expected in beta strands. The planarity of a putative amyloid core was represented by the highest average angle calculated. Models were only considered for experimental testing if this metric was below 20 degrees. Using FoldX 5.1 (52), all models were repaired with the RepairPDB command and had per-residue backbone hydrogen bond and total energies calculated with the SequenceDetail command. Putative amyloid cores with average total energy below 1 kcal/mol and average backbone hydrogen bond energy below -1 kcal/mol were considered for experimental testing. Structures were analyzed in Python 3.10.13 using pandas and NumPy. Data visualization was performed with matplotlib. Structure visualization was performed in R 4.3.1 with the r3dmol package.

### Yeast manipulations for DAmFRET and microscopy

Individual transformant colonies of rhy3279 were inoculated into 200μL standard synthetic media containing 2% dextrose (SD-Ura) in an Axygen® 96-well round-bottom assay plate. Untransformed yeast cells for color compensation were inoculated into nonselective media. Cells were incubated while shaking overnight on a Heidolph Titramax-1000 at 1000 rpm at 30°C. To induce ectopic protein expression and growth arrest in liquid media prior to cytometry or microscopy based analysis, cells were spun down and resuspended into 200μL in standard synthetic media containing 2% galactose (SGal-Ura) and 100μM MG132 (nonselective media for untransformed samples). Cells were induced and growth arrested for a total of 20 hr. 4 hr prior to analysis, cells were spun down and resuspended in fresh media and 100μm MG132. For samples analyzed via flow cytometry, photoconversion of samples was performed in a 96-well Axygen® assay plate while shaking at 1000 rpm. Cells were illuminated using an OmniCure S2000 Elite lamp fitted with a 320–500 nm (violet) filter and a beam collimator (Exfo), positioned 45 cm above the plate, for a duration of 5 min. Following photoconversion, cells were assayed using a BioRad ZE5 cell analyzer. To prevent sedimentation of cells, samples were periodically shaken for 5 sec during runtime. Donor fluorescence was excited with a 488 nm laser and collected from a 525/35 emission filter; FRET signal was excited with a 488 nm laser and collected from a 593/52 emission filter; Acceptor fluorescence was excited with a 561 nm laser and collected from a 589/15 emission filter; BDFP fluorescence was excited with a 640 nm laser and collected from a 670/30 emission filter; autofluorescence signal was excited with a 405 nm laser and collected from a 460/22 emission filter. Samples were manually compensated using untransformed cells as a fluorophore negative control, non-photoconverted mEos3.1 (donor), BDFP single color sample, and dsRed2 as a proxy for photoconverted mEos3.1 (acceptor).

### SDD-AGE

Transformed cells of rhy2145 and rhy3082 were inoculated, induced, harvested, lysed, and denatured as previously described using 0.1% sarkosyl ((6)). The gel was directly imaged using GE Typhoon Imaging System using a 488 laser and 525(40) BP filter. Images were then loaded into Fiji (2.14.0) for contrast adjustment.

### DAmFRET data analysis

Data were processed in FCS Express 7 7.22.0031 (De Novo Software). Events were selectively gated using forward scatter (FSC), side scatter (SSC), and the 460/22 emission channel (FSC 488/10-A, FSC 488/10-H, SSC 488/10-A, 460/22-405 nm-A) to include single, live, unbudded yeast cells, and exclude cellular debris and cells with high autofluorescence. All cells with donor fluorescence relative to the untransformed dark color control were included in final DAmFRET plots where the AmFRET parameter is the ratio of FRET intensity to Acceptor intensity, and concentration is the ratio of acceptor fluorescence to side scatter (SSC 488/10-A) as previously described (28).

To approximate C_sat_, a spline is fit to the mean AmFRET values across binned x-axis values. The spline was fit on binned values from the 1st the 99th percentile of values. The bins were further restricted to those with at least 20 cells, and a cell density of at least 150. The bin density is measured by the number of events divided by the interquartile range (IQR) of AmFRET values within the bin. The number of bins was determined using Scott’s rule for number of bins in a histogram. The mean AmFRET values were calculated for each bin, and then denoised using the Python SciPy.signal.sosfiltfilt, resulting in the spline fit. This was bootstrapped 100 times with the average and std of each bin of all splines reported as the final spline fit. The transition point is defined as the x-axis value that has the greatest change in AmFRET. Therefore, this value was calculated as the maximum of the first derivative of the fitted spline, using the numpy.diff package. The transition range is defined as the region between the maximum positive and negative rate of change, indicating where the transition between FRET states is starting and ending, respectively. The transition starting point is calculated as the maximum of the second derivative that lies before the transition point, and the transition endpoint is calculated as the minimum of the second derivative that lies after the transition point. This is done on all 100 bootstrapped splines and the median of bootstraps for three biological replicates is reported.

### Confocal microscopy

Confocal images of transformed yeast cells of lrhy3082 were acquired with an Andor Xyla x4 sCMOS camera at full resolution on a PerkinElmer Opera Phenix spinning disk microscope. Non-photoconverted mEos3.1 (green) was excited using a 488nm (20mw) laser at 20% for 20 ms and transmitted light was collected at 100% for 200 ms through a 40x water objective 1.1 NA (PerkinElmer). Emission was collected and filtered with 500-550nm (mEos green) and 650-760nm (transmitted) bandwidth emission filters. Z stacks were acquired at a z step spacing of 0.2μm for a total distance of 0.8μm. Each channel was max projected (NumPy) post-acquisition. Analysis was performed in Fiji version 2.14.0.

## Author contributions

**Hannah Kimbrough**: investigation; visualization; writing – original draft; writing – review and editing. **Jacob Jensen**: data curation; formal analysis; software; visualization. **Tayla Miller**: investigation, formal analysis. **Jeffrey J. Lange**: investigation and formal analysis. **Randal Halfmann**: conceptualization; funding acquisition; supervision; writing – original draft; writing – review and editing.

## Acknowledgments

We thank members of the Halfmann lab for constructive input, and members of the Microscopy and Cytometry cores of the Stowers Institute for assistance with experiments. This work was performed to fulfill, in part, requirements for HK’s thesis research in the Graduate School of the Stowers Institute for Medical Research. This work was supported by the National Institute of General Medical Sciences of the National Institutes of Health (Award Number R01GM130927, to RH) and the Stowers Institute for Medical Research. The funders had no role in study design, data collection and analysis, or manuscript preparation. The content is solely the responsibility of the authors and does not necessarily represent the official views of the funders.

## Supporting Information

Original data underlying this manuscript can be accessed from the Stowers Original Data Repository at https://www.stowers.org/research/publications/LIBPB-2522.

**Figure S1. Additional data supporting main figures**. (a) AlphaFold structural prediction of human functional amyloid Ripk3 (413-512). (b) AlphaFold pentamer predictions of all dipeptide repeat combinations at total peptide length 40 amino acids, colored by chain. (c). SDD-AGE gel prepared from [pin-] yeast cells expressing (TA)20, the intrinsically disordered domain of TDP43, and an amyloid forming polyQ tract where the dashed line represents the two adjacent groups of lanes are spliced from different positions in the gel. (d) Representative confocal microscopy image of yeast cells expressing (TTAA)10 at a range of intracellular concentrations. Scale bar, 10 μm. (e) AlphaFold structural predictions of (TA)20 (left) and (TTAA)10 (right) where backbone is colored by pLDDT and side chains are colored green (alanine) and grey (threonine).

**TableS1**. Plasmids used in this study.

