## Supplementary figures and images for "A tunable affinity fusion tag for protein self-assembly"

### Figure S1

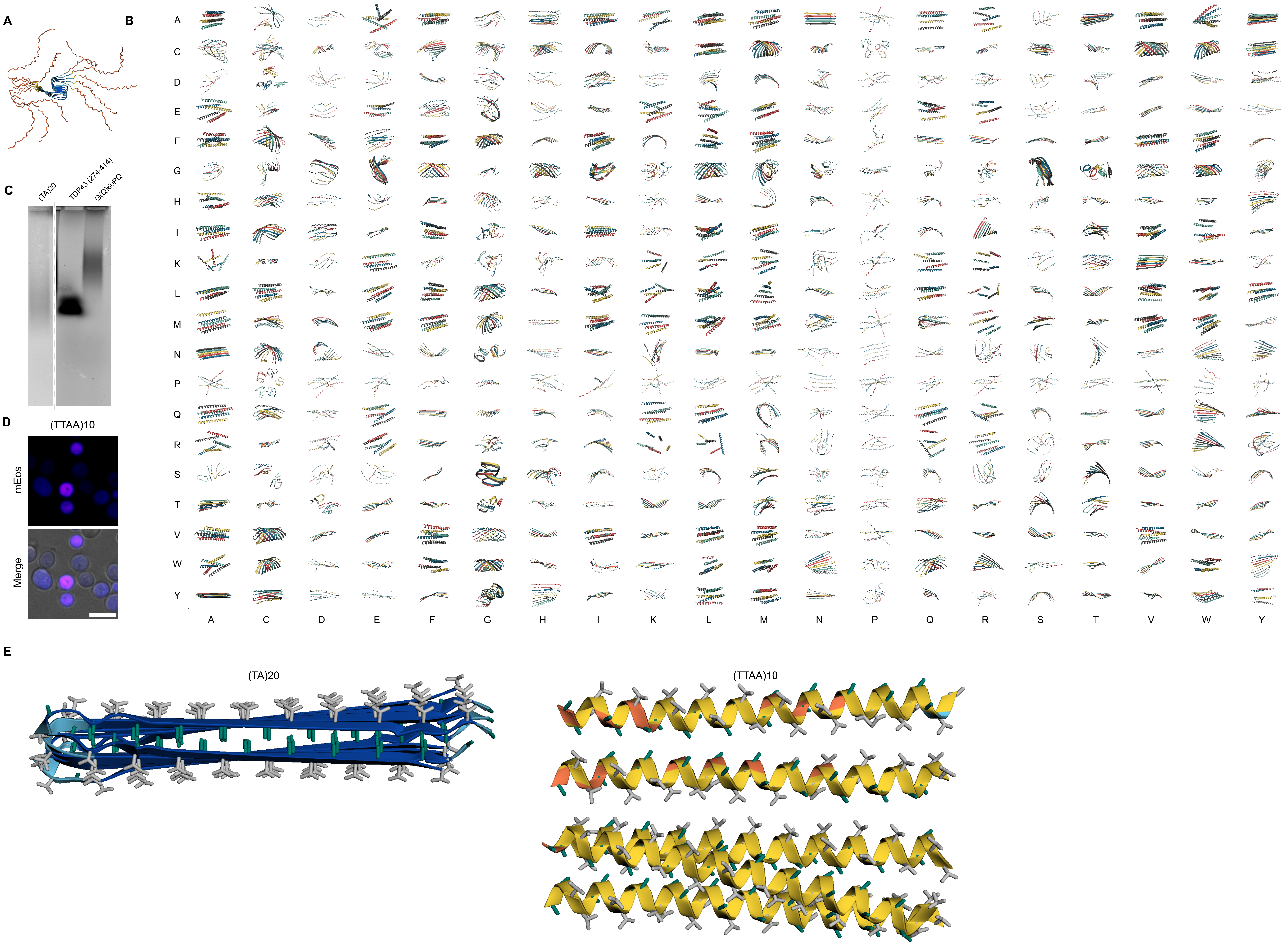
