## Supplementary material for "A tunable affinity fusion tag for protein self-assembly": Table S1

| Name | Insert | Source | Fusion protein sequence |
|---|---|---|---|
| rhx5639 | (YA)20 | This study | MSAIKPDMKIKLRMEGNVNGHHFVIDGDGTGKPFEGKQSMDLEVKEGGPLPFAFDILTTAFHYGNRVFAKYPDNIQDYFKQ<br>SFPKGYSWERSLTFEDGGICNARNDITMEGDTFYNKVRFYGTNFPANGPVMQKKTLKWEPSTEKMYVRDGVLTGDVEMALL<br>LEGNAHYRCDFRTTYKAKEKGVKLPGAHFVDHCIEILSHDKDYNKVKLYEHAVAHSGLPDNARRGAPEAAAREAAAREAAAR<br>EAAARGGAYAYAYAYAYAYAYAYAYAYAYAYAYAYAYAYAYAY |
| rhx3289 | (TA)20 | This study | MSAIKPDMKIKLRMEGNVNGHHFVIDGDGTGKPFEGKQSMDLEVKEGGPLPFAFDILTTAFHYGNRVFAKYPDNIQDYFKQ<br>SFPKGYSWERSLTFEDGGICNARNDITMEGDTFYNKVRFYGTNFPANGPVMQKKTLKWEPSTEKMYVRDGVLTGDVEMALL<br>LEGNAHYRCDFRTTYKAKEKGVKLPGAHFVDHCIEILSHDKDYNKVKLYEHAVAHSGLPDNARRGAPEAAAREAAAREAAAR<br>EAAARMSTATATATATATATATATATATATATATATATATATA |
| rhx5638 | (NA)20 | This study | MSAIKPDMKIKLRMEGNVNGHHFVIDGDGTGKPFEGKQSMDLEVKEGGPLPFAFDILTTAFHYGNRVFAKYPDNIQDYFKQ<br>SFPKGYSWERSLTFEDGGICNARNDITMEGDTFYNKVRFYGTNFPANGPVMQKKTLKWEPSTEKMYVRDGVLTGDVEMALL<br>LEGNAHYRCDFRTTYKAKEKGVKLPGAHFVDHCIEILSHDKDYNKVKLYEHAVAHSGLPDNARRGAPEAAAREAAAREAAAR<br>EAAARGGANANANANANANANANANANANANANANANANANAN |
| rhx4299 | (GA)20 | This study | MSAIKPDMKIKLRMEGNVNGHHFVIDGDGTGKPFEGKQSMDLEVKEGGPLPFAFDILTTAFHYGNRVFAKYPDNIQDYFKQ<br>SFPKGYSWERSLTFEDGGICNARNDITMEGDTFYNKVRFYGTNFPANGPVMQKKTLKWEPSTEKMYVRDGVLTGDVEMALL<br>LEGNAHYRCDFRTTYKAKEKGVKLPGAHFVDHCIEILSHDKDYNKVKLYEHAVAHSGLPDNARRGAPEAAAREAAAREAAAR<br>EAAARMSGAGAGAGAGAGAGAGAGAGAGAGAGAGAGAGAGAGA |
| rhx5641 | (YA)10 | This study | MSAIKPDMKIKLRMEGNVNGHHFVIDGDGTGKPFEGKQSMDLEVKEGGPLPFAFDILTTAFHYGNRVFAKYPDNIQDYFKQ<br>SFPKGYSWERSLTFEDGGICNARNDITMEGDTFYNKVRFYGTNFPANGPVMQKKTLKWEPSTEKMYVRDGVLTGDVEMALL<br>LEGNAHYRCDFRTTYKAKEKGVKLPGAHFVDHCIEILSHDKDYNKVKLYEHAVAHSGLPDNARRGAPEAAAREAAAREAAAR<br>EAAARGGAYAYAYAYAYAYAYAYAYAYAY |
| rhx3290 | (TA)10 | This study | MSAIKPDMKIKLRMEGNVNGHHFVIDGDGTGKPFEGKQSMDLEVKEGGPLPFAFDILTTAFHYGNRVFAKYPDNIQDYFKQ<br>SFPKGYSWERSLTFEDGGICNARNDITMEGDTFYNKVRFYGTNFPANGPVMQKKTLKWEPSTEKMYVRDGVLTGDVEMALL<br>LEGNAHYRCDFRTTYKAKEKGVKLPGAHFVDHCIEILSHDKDYNKVKLYEHAVAHSGLPDNARRGAPEAAAREAAAREAAAR<br>EAAARMSTATATATATATATATATATATA |
| rhx3864 | (GA)60 | This study | MSAIKPDMKIKLRMEGNVNGHHFVIDGDGTGKPFEGKQSMDLEVKEGGPLPFAFDILTTAFHYGNRVFAKYPDNIQDYFKQ<br>SFPKGYSWERSLTFEDGGICNARNDITMEGDTFYNKVRFYGTNFPANGPVMQKKTLKWEPSTEKMYVRDGVLTGDVEMALL<br>LEGNAHYRCDFRTTYKAKEKGVKLPGAHFVDHCIEILSHDKDYNKVKLYEHAVAHSGLPDNARRGAPEAAAREAAAREAAAR<br>EAAARMSGAGAGAGAGAGAGAGAGAGAGAGAGAGAGAGAGAGAGAGAGAGAGAGAGAGAGAGAGAGAGAGAGAGAGA<br>GAGAGAGAGAGAGAGAGAGAGAGAGAGAGAGAGAGAGAGAGAGAGA |
| rhx4101 | (TA)8 | This study | MSAIKPDMKIKLRMEGNVNGHHFVIDGDGTGKPFEGKQSMDLEVKEGGPLPFAFDILTTAFHYGNRVFAKYPDNIQDYFKQ<br>SFPKGYSWERSLTFEDGGICNARNDITMEGDTFYNKVRFYGTNFPANGPVMQKKTLKWEPSTEKMYVRDGVLTGDVEMALL<br>LEGNAHYRCDFRTTYKAKEKGVKLPGAHFVDHCIEILSHDKDYNKVKLYEHAVAHSGLPDNARRGAPEAAAREAAAREAAAR<br>EAAARMSTATATATATATATATA |
| rhx3444 | (TA)12 | This study | MSAIKPDMKIKLRMEGNVNGHHFVIDGDGTGKPFEGKQSMDLEVKEGGPLPFAFDILTTAFHYGNRVFAKYPDNIQDYFKQ<br>SFPKGYSWERSLTFEDGGICNARNDITMEGDTFYNKVRFYGTNFPANGPVMQKKTLKWEPSTEKMYVRDGVLTGDVEMALL<br>LEGNAHYRCDFRTTYKAKEKGVKLPGAHFVDHCIEILSHDKDYNKVKLYEHAVAHSGLPDNARRGAPEAAAREAAAREAAAR<br>EAAARMSTATATATATATATATATATATATA |
| rhx3443 | (TA)14 | This study | MSAIKPDMKIKLRMEGNVNGHHFVIDGDGTGKPFEGKQSMDLEVKEGGPLPFAFDILTTAFHYGNRVFAKYPDNIQDYFKQ<br>SFPKGYSWERSLTFEDGGICNARNDITMEGDTFYNKVRFYGTNFPANGPVMQKKTLKWEPSTEKMYVRDGVLTGDVEMALL<br>LEGNAHYRCDFRTTYKAKEKGVKLPGAHFVDHCIEILSHDKDYNKVKLYEHAVAHSGLPDNARRGAPEAAAREAAAREAAAR<br>EAAARMSTATATATATATATATATATATATATATA |
| rhx3456 | (TTAA)10 | This study | MSAIKPDMKIKLRMEGNVNGHHFVIDGDGTGKPFEGKQSMDLEVKEGGPLPFAFDILTTAFHYGNRVFAKYPDNIQDYFKQ<br>SFPKGYSWERSLTFEDGGICNARNDITMEGDTFYNKVRFYGTNFPANGPVMQKKTLKWEPSTEKMYVRDGVLTGDVEMALL<br>LEGNAHYRCDFRTTYKAKEKGVKLPGAHFVDHCIEILSHDKDYNKVKLYEHAVAHSGLPDNARRGAPEAAAREAAAREAAAR<br>EAAARMSTTAATTAATTAATTAATTAATTAATTAATTAATTAATTAA |
| rhx5429 | 8\|8 | This study | MSAIKPDMKIKLRMEGNVNGHHFVIDGDGTGKPFEGKQSMDLEVKEGGPLPFAFDILTTAFHYGNRVFAKYPDNIQDYFKQ<br>SFPKGYSWERSLTFEDGGICNARNDITMEGDTFYNKVRFYGTNFPANGPVMQKKTLKWEPSTEKMYVRDGVLTGDVEMALL<br>LEGNAHYRCDFRTTYKAKEKGVKLPGAHFVDHCIEILSHDKDYNKVKLYEHAVAHSGLPDNARRGAPEAAAREAAAREAAAR<br>EAAARMSTAATTAATTAATATATATATTAATATATATAT |
| rhx5428 | 8\|9 | This study | MSAIKPDMKIKLRMEGNVNGHHFVIDGDGTGKPFEGKQSMDLEVKEGGPLPFAFDILTTAFHYGNRVFAKYPDNIQDYFKQ<br>SFPKGYSWERSLTFEDGGICNARNDITMEGDTFYNKVRFYGTNFPANGPVMQKKTLKWEPSTEKMYVRDGVLTGDVEMALL<br>LEGNAHYRCDFRTTYKAKEKGVKLPGAHFVDHCIEILSHDKDYNKVKLYEHAVAHSGLPDNARRGAPEAAAREAAAREAAAR<br>EAAARMSTATTAATTAATATATATATTAATATATATA |
| rhx5427 | 9\|9 | This study | MSAIKPDMKIKLRMEGNVNGHHFVIDGDGTGKPFEGKQSMDLEVKEGGPLPFAFDILTTAFHYGNRVFAKYPDNIQDYFKQ<br>SFPKGYSWERSLTFEDGGICNARNDITMEGDTFYNKVRFYGTNFPANGPVMQKKTLKWEPSTEKMYVRDGVLTGDVEMALL<br>LEGNAHYRCDFRTTYKAKEKGVKLPGAHFVDHCIEILSHDKDYNKVKLYEHAVAHSGLPDNARRGAPEAAAREAAAREAAAR<br>EAAARMSTAATTAATTATATATATATTAATATATATA |
| rhx5426 | 9\|10 | This study | MSAIKPDMKIKLRMEGNVNGHHFVIDGDGTGKPFEGKQSMDLEVKEGGPLPFAFDILTTAFHYGNRVFAKYPDNIQDYFKQ<br>SFPKGYSWERSLTFEDGGICNARNDITMEGDTFYNKVRFYGTNFPANGPVMQKKTLKWEPSTEKMYVRDGVLTGDVEMALL<br>LEGNAHYRCDFRTTYKAKEKGVKLPGAHFVDHCIEILSHDKDYNKVKLYEHAVAHSGLPDNARRGAPEAAAREAAAREAAAR<br>EAAARMSTAATAATTATATATATATTAATATATATAT |
| rhx5425 | 10\|10 | This study | MSAIKPDMKIKLRMEGNVNGHHFVIDGDGTGKPFEGKQSMDLEVKEGGPLPFAFDILTTAFHYGNRVFAKYPDNIQDYFKQ<br>SFPKGYSWERSLTFEDGGICNARNDITMEGDTFYNKVRFYGTNFPANGPVMQKKTLKWEPSTEKMYVRDGVLTGDVEMALL<br>LEGNAHYRCDFRTTYKAKEKGVKLPGAHFVDHCIEILSHDKDYNKVKLYEHAVAHSGLPDNARRGAPEAAAREAAAREAAAR<br>EAAARMSTAATTAATATATATATATTAATATATATAT |
| rhx5424 | 10\|11 | This study | MSAIKPDMKIKLRMEGNVNGHHFVIDGDGTGKPFEGKQSMDLEVKEGGPLPFAFDILTTAFHYGNRVFAKYPDNIQDYFKQ<br>SFPKGYSWERSLTFEDGGICNARNDITMEGDTFYNKVRFYGTNFPANGPVMQKKTLKWEPSTEKMYVRDGVLTGDVEMALL<br>LEGNAHYRCDFRTTYKAKEKGVKLPGAHFVDHCIEILSHDKDYNKVKLYEHAVAHSGLPDNARRGAPEAAAREAAAREAAAR<br>EAAARMSTATTAATATATATATATTAATATATATATA |
| rhx5423 | 11\|11 | This study | MSAIKPDMKIKLRMEGNVNGHHFVIDGDGTGKPFEGKQSMDLEVKEGGPLPFAFDILTTAFHYGNRVFAKYPDNIQDYFKQ<br>SFPKGYSWERSLTFEDGGICNARNDITMEGDTFYNKVRFYGTNFPANGPVMQKKTLKWEPSTEKMYVRDGVLTGDVEMALL<br>LEGNAHYRCDFRTTYKAKEKGVKLPGAHFVDHCIEILSHDKDYNKVKLYEHAVAHSGLPDNARRGAPEAAAREAAAREAAAR<br>EAAARMSTAATTATATATATATATTAATATATATATA |
| rhx5380 | 12\|11 | This study | MSAIKPDMKIKLRMEGNVNGHHFVIDGDGTGKPFEGKQSMDLEVKEGGPLPFAFDILTTAFHYGNRVFAKYPDNIQDYFKQ<br>SFPKGYSWERSLTFEDGGICNARNDITMEGDTFYNKVRFYGTNFPANGPVMQKKTLKWEPSTEKMYVRDGVLTGDVEMALL<br>LEGNAHYRCDFRTTYKAKEKGVKLPGAHFVDHCIEILSHDKDYNKVKLYEHAVAHSGLPDNARRGAPEAAAREAAAREAAAR<br>EAAARMSTTAATATATATATATATTAATATATATATA |
| rhx5422 | 12\|12 | This study | MSAIKPDMKIKLRMEGNVNGHHFVIDGDGTGKPFEGKQSMDLEVKEGGPLPFAFDILTTAFHYGNRVFAKYPDNIQDYFKQ<br>SFPKGYSWERSLTFEDGGICNARNDITMEGDTFYNKVRFYGTNFPANGPVMQKKTLKWEPSTEKMYVRDGVLTGDVEMALL<br>LEGNAHYRCDFRTTYKAKEKGVKLPGAHFVDHCIEILSHDKDYNKVKLYEHAVAHSGLPDNARRGAPEAAAREAAAREAAAR<br>EAAARMSTAATATATATATATATTAATATATATATAT |
| rhx5381 | 13\|13 | This study | MSAIKPDMKIKLRMEGNVNGHHFVIDGDGTGKPFEGKQSMDLEVKEGGPLPFAFDILTTAFHYGNRVFAKYPDNIQDYFKQ<br>SFPKGYSWERSLTFEDGGICNARNDITMEGDTFYNKVRFYGTNFPANGPVMQKKTLKWEPSTEKMYVRDGVLTGDVEMALL<br>LEGNAHYRCDFRTTYKAKEKGVKLPGAHFVDHCIEILSHDKDYNKVKLYEHAVAHSGLPDNARRGAPEAAAREAAAREAAAR<br>EAAARMSTATATATATATATATTAATATATATATATA |
| rhx1900b | TDP-43 274-414 | This study | MSAIKPDMKIKLRMEGNVNGHHFVIDGDGTGKPFEGKQSMDLEVKEGGPLPFAFDILTTAFHYGNRVFAKYPDNIQDYFKQ<br>SFPKGYSWERSLTFEDGGICNARNDITMEGDTFYNKVRFYGTNFPANGPVMQKKTLKWEPSTEKMYVRDGVLTGDVEMALL<br>LEGNAHYRCDFRTTYKAKEKGVKLPGAHFVDHCIEILSHDKDYNKVKLYEHAVAHSGLPDNARRGAPEAAAREAAAREAAAR<br>EAAARMSGRFGGNPGGFGNQGGFGNSRGGGAGLGNNQGSNMGGGMNFGAFSINPAMMAAAQAALQSSWGMMGML<br>ASQQNQSGPSGNNQNQGNMQREPNQAFGSGNNSYSGSNSGAAIGWGSASNAGSGSGFNGGFGSSMDSKSSGWGM |
| rhx2682 | (Q)60 | Kandola et al. 2023 | |
| rhx5530 | MBP-12\|12 | This study | MSKIEEGKLVIWINGDKGYNGLAEVGKKFEKDTGIKVTVEHPDKLEEKFPQVAATGDGPDIIFWAHDRFGGYAQSGLLAEITP<br>DKAFQDKLYPFTWDAVRYNGKLIAYPIAVEALSLIYNKDLLPNPPKTWEEIPALDKELKAKGKSALMFNLQEPYFTWPLIAADG<br>GYAFKYENGKYDIKDVGVDNAGAKAGLTFLVDLIKNKHMNADTDYSIAEAAFNKGETAMTINGPWAWSNIDTSKVNYGVTVL<br>PTFKGQPSKPFVGVLSAGINAASPNKELAKEFLENYLLTDEGLEAVNKDKPLGAVALKSYEEELAKDPRIAATMENAQKGEIMP<br>NIPQMSAFWYAVRTAVINAASGRQTVDEALKDAQTGGGSAIKPDMKIKLRMEGNVNGHHFVIDGDGTGKPFEGKQSMDLE<br>VKEGGPLPFAFDILTTAFHYGNRVFAKYPDNIQDYFKQSFPKGYSWERSLTFEDGGICNARNDITMEGDTFYNKVRFYGTNFP<br>ANGPVMQKKTLKWEPSTEKMYVRDGVLTGDVEMALLLEGNAHYRCDFRTTYKAKEKGVKLPGAHFVDHCIEILSHDKDYNK<br>VKLYEHAVAHSGLPDNARRGAPEAAAREAAAREAAAREAAARMSTAATATATATATATTAATATATATATAT |
| rhx5531 | MBP-10\|10 | This study | MSKIEEGKLVIWINGDKGYNGLAEVGKKFEKDTGIKVTVEHPDKLEEKFPQVAATGDGPDIIFWAHDRFGGYAQSGLLAEITP<br>DKAFQDKLYPFTWDAVRYNGKLIAYPIAVEALSLIYNKDLLPNPPKTWEEIPALDKELKAKGKSALMFNLQEPYFTWPLIAADG<br>GYAFKYENGKYDIKDVGVDNAGAKAGLTFLVDLIKNKHMNADTDYSIAEAAFNKGETAMTINGPWAWSNIDTSKVNYGVTVL<br>PTFKGQPSKPFVGVLSAGINAASPNKELAKEFLENYLLTDEGLEAVNKDKPLGAVALKSYEEELAKDPRIAATMENAQKGEIMP<br>NIPQMSAFWYAVRTAVINAASGRQTVDEALKDAQTGGGSAIKPDMKIKLRMEGNVNGHHFVIDGDGTGKPFEGKQSMDLE<br>VKEGGPLPFAFDILTTAFHYGNRVFAKYPDNIQDYFKQSFPKGYSWERSLTFEDGGICNARNDITMEGDTFYNKVRFYGTNFP<br>ANGPVMQKKTLKWEPSTEKMYVRDGVLTGDVEMALLLEGNAHYRCDFRTTYKAKEKGVKLPGAHFVDHCIEILSHDKDYNK<br>VKLYEHAVAHSGLPDNARRGAPEAAAREAAAREAAAREAAARMSTAATTAATATATATATTAATATATATAT |
| rhx5532 | EB250-12\|12 | This study | MSEEQPEPIVSEQEDPEEPGSSVFEILPPEQSPGGPPEEPDQPSEEPVMEEIQPQLEGSPGPPPEGPPEPDPEEDESEEIQQ<br>PISQPSPPSEGQLLEPLQEQSSSSEESGPSEPSSDPSSEESDPPEPLISVFPSEGSSEPPQQPDDLSPPLEPQPEEQPEPGSP<br>DDQSPPPSPSPPGEPQGQPDGSPSGEPGQSEEPQPGGDPEIVPPIQEELPEPSPEGSPLEGSIGEMVSPPGSDPESDGGP<br>QPPQDQQAIKPDMKIKLRMEGNVNGHHFVIDGDGTGKPFEGKQSMDLEVKEGGPLPFAFDILTTAFHYGNRVFAKYPDNIQ<br>DYFKQSFPKGYSWERSLTFEDGGICNARNDITMEGDTFYNKVRFYGTNFPANGPVMQKKTLKWEPSTEKMYVRDGVLTGDV<br>EMALLLEGNAHYRCDFRTTYKAKEKGVKLPGAHFVDHCIEILSHDKDYNKVKLYEHAVAHSGLPDNARRGAPEAAAREAAAR<br>EAAAREAAARMSTAATATATATATATTAATATATATATAT |
| rhx5533 | EB250-10\|10 | This study | MSEEQPEPIVSEQEDPEEPGSSVFEILPPEQSPGGPPEEPDQPSEEPVMEEIQPQLEGSPGPPPEGPPEPDPEEDESEEIQQ<br>PISQPSPPSEGQLLEPLQEQSSSSEESGPSEPSSDPSSEESDPPEPLISVFPSEGSSEPPQQPDDLSPPLEPQPEEQPEPGSP<br>DDQSPPPSPSPPGEPQGQPDGSPSGEPGQSEEPQPGGDPEIVPPIQEELPEPSPEGSPLEGSIGEMVSPPGSDPESDGGP<br>QPPQDQQAIKPDMKIKLRMEGNVNGHHFVIDGDGTGKPFEGKQSMDLEVKEGGPLPFAFDILTTAFHYGNRVFAKYPDNIQ<br>DYFKQSFPKGYSWERSLTFEDGGICNARNDITMEGDTFYNKVRFYGTNFPANGPVMQKKTLKWEPSTEKMYVRDGVLTGDV<br>EMALLLEGNAHYRCDFRTTYKAKEKGVKLPGAHFVDHCIEILSHDKDYNKVKLYEHAVAHSGLPDNARRGAPEAAAREAAAR<br>EAAAREAAARMSTAATTAATATATATATTAATATATATAT |
